# Bioorthogonal labeling enables in situ fluorescence imaging of expressed gas vesicle nanostructures

**DOI:** 10.1101/2023.11.30.569486

**Authors:** Erik Schrunk, Przemysław Dutka, Robert C. Hurt, Di Wu, Mikhail G. Shapiro

## Abstract

Gas vesicles (GVs) are proteinaceous nanostructures that, along with virus-like particles, encapsulins, nano-cages, and other macromolecular assemblies are being developed for potential biomedical applications. To facilitate such development, it would be valuable to characterize these nanostructures’ sub-cellular assembly and localization. However, traditional fluorescent protein fusions are not tolerated by GVs’ primary constituent protein, making optical microscopy a challenge. Here, we introduce a method for fluorescently visualizing intracellular GVs using the bioorthogonal label FlAsH, which becomes fluorescent upon binding the six-amino acid tetracysteine (TC) tag. We engineered the GV subunit protein, GvpA, to display the TC tag, and showed that GVs bearing TC-tagged GvpA can be successfully assembled and fluorescently visualized in HEK 293T cells. We used fluorescence images of the tagged GVs to study GV size and distance distributions within these cells. This bioorthogonal labeling approach will enable research to provide a greater understanding of GVs and could be adapted to similar proteinaceous nanostructures.

## INTRODUCTION

Gas vesicles (GVs) are air-filled protein nanostructures (∼85 nm diameter, ∼500 nm length)^1^ that are entering use in biomedical applications alongside other proteinaceous macromolecular assemblies such as encapsulins, virus-like particles and nano-cages.^2–4^ In particular, GVs have recently emerged as promising agents for biomolecular ultrasound: they have been expressed recombinantly in both bacterial and mammalian cells and have been used as cavitation nuclei,^5^ ultrasonic reporters of cancer,^6^ acoustic actuators for selective cellular manipulation,^7^ and more.^8–11^ These GV-based technologies—and those involving other macromolecular complexes—could benefit from knowledge of these structures’ sub-cellular localization, as this knowledge could enable the engineering of systems targeted to specific organelles or cellular compartments. However, there are currently no reported methods to fluorescently label GVs within cells. In part, this is because the composition of GVs as assemblies of small, highly conserved subunit proteins makes it difficult for them to accommodate substantial fused functionalities such as fluorescent proteins.

Here, we describe a method to optically visualize GVs inside cells by genetically modifying the GV shell protein GvpA with the tetracysteine (TC) motif, allowing the GVs to be fluorescently labeled with the bioorthogonal FlAsH reagent for visualization of their sub-cellular localization. FlAsH is a membrane-permeable fluorogenic molecule that binds specifically with the TC tag (Cys-Cys-Pro-Gly-Cys-Cys) and which turns on fluorescence upon binding.^12–15^ We sought to introduce this tag into the major GV structural protein, GvpA, such that expressed intracellular GVs would be able to bind FlAsH and turn on fluorescence. We screened for TC-containing GvpA mutants in bacteria and, once we identified a suitable variant, expressed TC-tagged *Anabaena flosaquae* GVs (“AnaGVs”) in HEK 293T cells. Using these “tcGVs,” we were able to directly visualize the three-dimensional distribution of GVs in the cell, observing that they tend to form clusters in the cytosol. In addition to enabling the study of GVs, this approach will inform similar studies of other genetically encoded protein nanostructures.

## RESULTS AND DISCUSSION

### The C-terminus of GvpA is amenable to single substitutions to cysteine

To engineer tcGVs, we sought to incorporate the TC motif into the GV shell protein GvpA. We looked to introduce the motif into a region of GvpA which faces the GV exterior—thereby making it accessible to cytosolic FlAsH—and which is tolerant of mutations to cysteine, such that introduction of the TC tag does not abrogate GvpA expression and GV assembly. To predict which region of GvpA would best accommodate the TC tag, we looked to structural models of GvpA.^1,16^ In a random mutagenesis experiment, the C-terminus of GvpA was found to be tolerant of many different point mutations – more so than any other region of the protein – suggesting that this region could be the most amenable to the substitution of the six-amino acid TC tag.^1^ Furthermore, the C-terminus of GvpA is on the exterior-facing region of the protein^1^ (Figure 1a-c). We therefore selected the C-terminus of GvpA as our target location.

**Figure 1:**
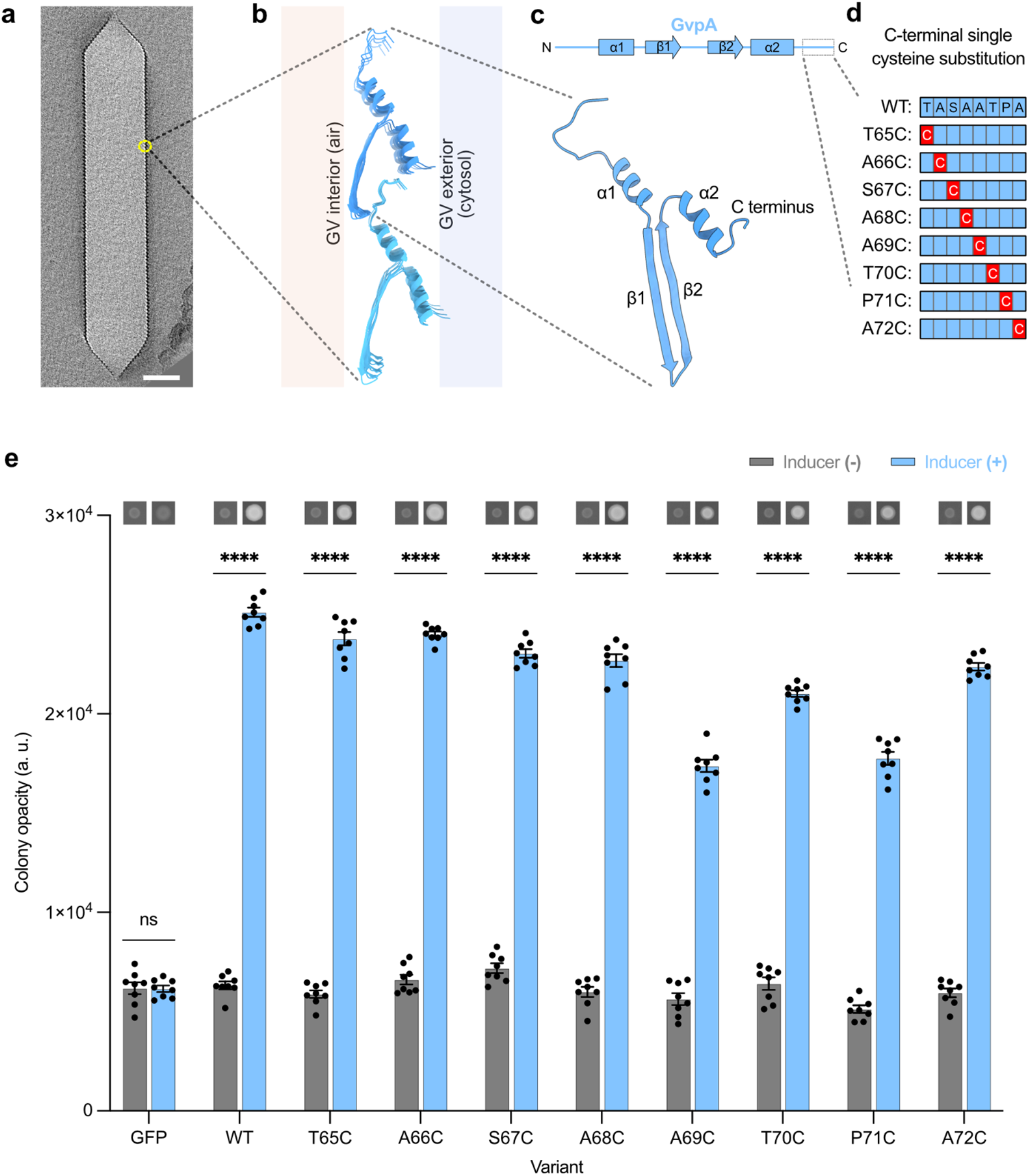
The C-terminus of the gas vesicle structural protein GvpA is tolerant of point mutations to cysteine. (a) A central slice through an electron cryo-tomogram of an individual AnaGV (EMD-29922)^1^. Scale bar 50 nm. (b) Atomic model of two adjacent “ribs” composed of GvpA (PDB 8GBS)^1^. Interior- and exterior-facing sides of the GV shell indicated. (c) Linear and atomic models of a single GvpA molecule with alpha helix and beta strand regions indicated.^1^ The C-terminal region of GvpA is shown by a dashed box in the linear representation of GvpA. (d) Schematic of the C-terminal variants of GvpA1 screened. Each red box represents a point mutation to cysteine and each blue box represents the amino acid in the wild type GvpA1. (e) Graph of opacity for induced and uninduced bacterial patches transformed with plasmids coding for mutant GV expression. Colony opacity is indicative of GV expression. Representative images of induced and uninduced patches displayed above their corresponding columns in the graph. N = 8 patches per condition. Patches with a plasmid encoding green fluorescent protein (GFP) expression included as a GV-negative control. Asterisks represent statistical significance by unpaired *t*-tests (****: p<0.0001, ns: not significant). Error bars represent mean ± SEM.

Before attempting the substitution of four cysteines into GvpA, we first tested the ability of individual positions within its C-terminus to accommodate single-Cys mutations. We screened mutants in *E. coli* using the bARG_Ser_ construct, which uses GV genes derived from *Serratia sp*. 39006, including the GvpA homolog GvpA1^6^ (∼92% similarity to GvpA, sequence alignment in Figure S1). We mutated each of the final eight amino acids in GvpA1 to Cys (Figure 1d-e), then expressed the mutant GVs in confluent bacterial patches on Petri dishes containing the inducer arabinose. We then measured the opacity of the patches as a proxy for GV expression, as GVs scatter visible light.^17–19^ We observed GV expression in all mutants, with only modest reductions at positions 69-71 (Figure 1e), and concluded that the C-terminus of GvpA1 could tolerate point mutations to Cys.

### The C-terminus of GvpA is amenable to substitutions to the TC tag

With the knowledge that each amino acid in the C-terminus of GvpA1 could be individually substituted to Cys, we next tested multi-position substitutions to introduce the TC tag. We cloned three variants of the GvpA1 gene with the minimal TC tag (Cys-Cys-Xxx-Xxx-Cys-Cys) in all three possible C-terminal positions (Figure 2a), leaving the middle two non-Cys amino acids of the tag unchanged relative to wild type (WT) GvpA1 (denoted by Xxx) to minimize sequence disruption. We found GV expression in all cases (Figure 2b), although at reduced levels compared to wild type. The variant with the TC tag at the most C-terminal position, called TC3, had the highest opacity (Figure 2b) and the healthiest patch morphology (Figure S2), suggesting that this variant was the best tolerated by cells expressing the resulting GVs. As an additional test, we converted the two non-Cys residues in TC3 to Pro-Gly to create a full TC tag (Cys-Cys-Pro-Gly-Cys-Cys) (Figure 2a) and noted that this mutant, called TC4, expressed GVs as well (Figure 2b). We concluded that the optimal positioning of the TC tag in the C-terminus of GvpA1 was at the most C-terminal position.

**Figure 2:**
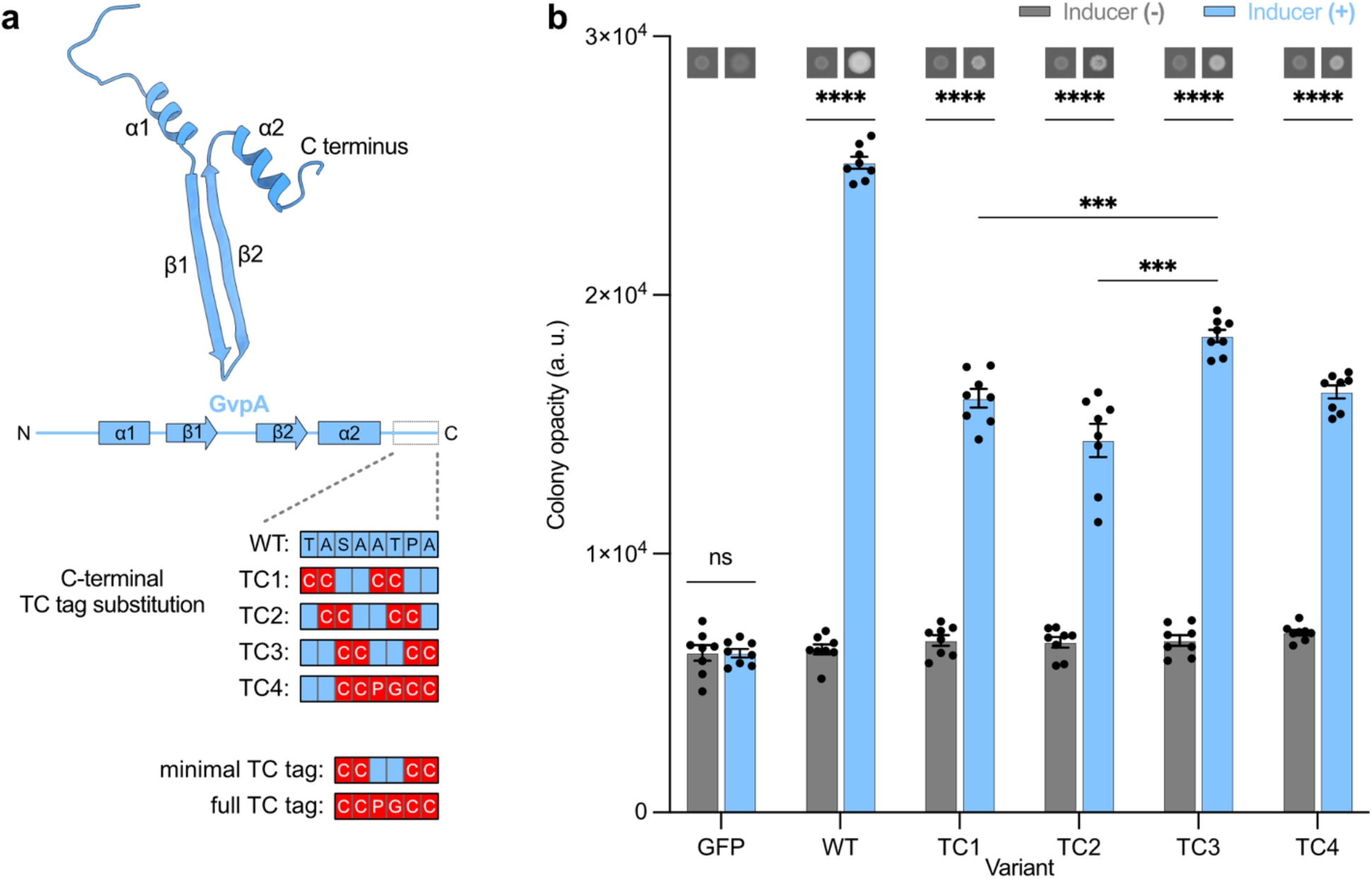
The C-terminus of GvpA tolerates substitution to the tetracysteine motifs CCXXCC and CCPGCC. (a) Schematic of the C-terminal tetracysteine mutants of GvpA1 screened. Each red “C” box represents a point mutation to cysteine and each blue box represents the amino acid in wild type GvpA1. TC1 through TC3 are the minimal TC tag Cys-Cys-Xxx-Xxx-Cys-Cys, while TC4 is the full TC tag Cys-Cys-Pro-Gly-Cys-Cys. (b) Graph of opacity for induced and uninduced bacterial patches transformed with plasmids coding for mutant GV expression. Colony opacity is indicative of GV expression. Representative images of induced and uninduced patches displayed above their corresponding columns in the graph. N = 8 patches per condition. Patches with a plasmid encoding GFP expression included as a GV-negative control. Asterisks represent statistical significance by unpaired *t*-tests (****: p<0.0001, ***: p<0.001, ns: not significant). Error bars represent mean ± SEM.

### Tetracysteine-tagged GvpA can be incorporated into GVs expressed in mammalian cells and imaged fluorescently by FlAsH

After establishing tcGV expression in bacteria, we set out to translate our approach to mammalian cells. We inserted the TC tag into the GvpA of the mARG construct^6^ at the same location as the best-performing tetracysteine-tagged GvpA1 from our bacterial screen (TC4) and called the resulting gene “tcGvpA.” We transfected human HEK 293T cells with mARG, replacing 10%, 20%, 25%, 50%, and 100% of the wild type GvpA (“wtGvpA”) with the tcGvpA in the transfection mixture, and observed GV formation in all but the 100% tcGvpA condition (Figure S3). This shows that while some wtGvpA is necessary for GV formation, tcGvpA expression is well-tolerated by mammalian cells. To determine whether tcGvpA is incorporated into the GVs, we next treated the transfected (20% tcGvpA) cells with FlAsH and found that FlAsH readily labeled the GVs in those cells (Figure 3b). Control cells expressing wild type GVs (“wtGVs”) without tcGvpA did not show labeling (Figure 3b). This demonstrated that tcGvpA is incorporated into mammalian GVs when co-expressed with wtGvpA and that the resulting chimeric GVs can be labeled intracellularly by FlAsH.

**Figure 3:**
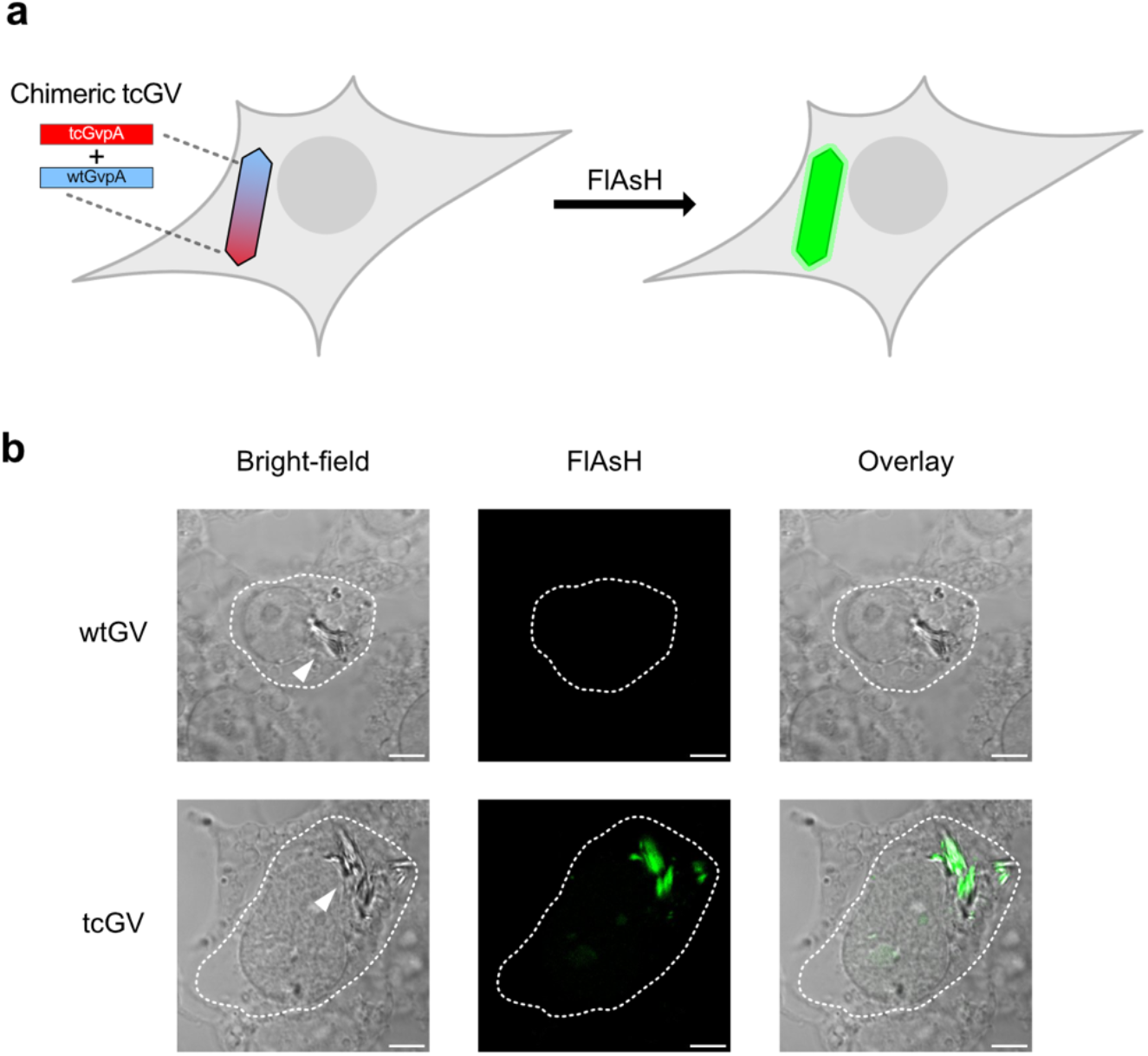
tcGVs can be successfully expressed and labeled with FlAsH in HEK 293T cells. (a) Schematic of an expressed tcGV cluster becoming fluorescent with FlAsH. tcGVs are comprised of wtGvpA and tcGvpA. After addition of FlAsH, the tcGVs become fluorescent as FlAsH binds to tcGvpA. (b) Images of wtGV and tcGV (20% tcGvpA) clusters in fixed HEK 293T cells (indicated by arrows). All GVs are visible under bright-field imaging (first column), but only tcGVs have any FlAsH signal above the background (second column). The bright-field/FlAsH overlay (third column) demonstrates that the strongest FlAsH signal overlaps with tcGV clusters. All scale bars 5 μm. GV-expressing cells outlined in white.

The co-delivery of tcGvpA and wtGvpA for tcGV expression is an important aspect of these findings, as it demonstrates that not every protein subunit of the GVs needs to be TC-tagged for the GV itself to be sufficiently reactive towards FlAsH. Therefore, the use of a mixture of wtGvpA and tcGvpA—notably with a significant majority of the wtGvpA gene—to express tcGVs highlights the utility of this approach when attempting to fluorescently label proteinaceous nanostructures within cells: even if TC-tagging a protein subunit is not ideal, spiking in a small fraction of TC-tagged subunits can be sufficient for informative labeling.

### GVs expressed in HEK 293T cells form clusters in the cytosol

After demonstrating that intracellular tcGVs could be fluorescently labeled with FlAsH, we sought to determine their subcellular location in mammalian cells. Knowledge of the localization of GVs within cells could improve our understanding of the biosynthesis and degradation of these protein structures and inform efforts to target GVs to specific organelles or cellular structures. Although phase contrast microscopy can be used to observe the presence of GVs within cells due to their differential refractive index,^19^ it does not provide reliable information about their subcellular localization due to poor depth resolution. On the other hand, imaging the GVs using confocal microscopy, now enabled by FlAsH labeling, would allow the determination of their precise subcellular location in 3D. To demonstrate this capability, we acquired multiple horizontal planes of cells expressing tcGVs labeled with FlAsH and simultaneously stained the nucleus with DAPI and the plasma membrane with a membrane-trafficked fluorescent protein^20^ (Lck-mScarlet-I) (Figure 4a; 3D renderings of additional cells are in Figure S4).

**Figure 4:**
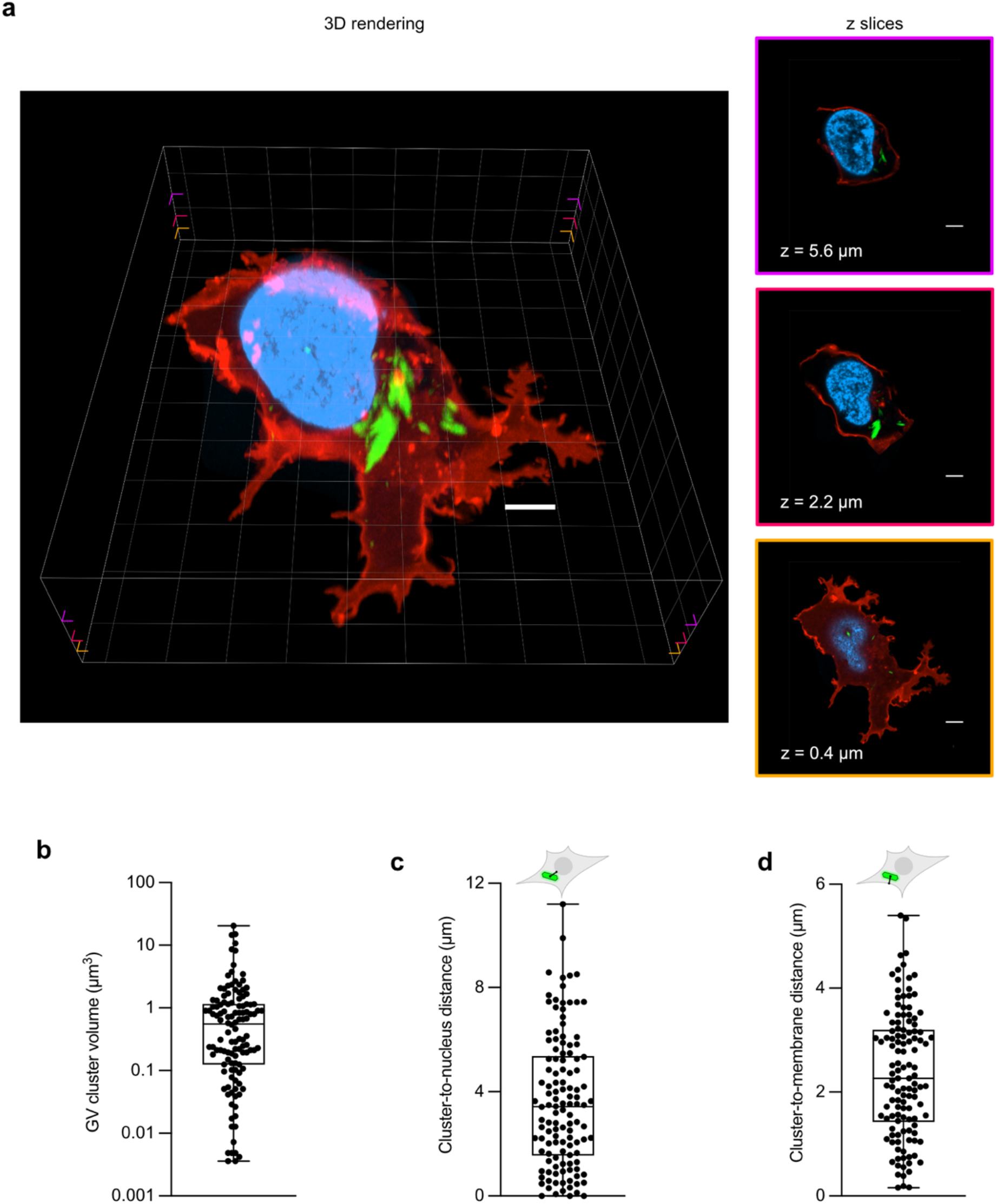
Fluorescence imaging of tcGVs elucidates the size and spatial distributions of GV clusters in HEK 293T cells. (a) A 3D rendering (left) of a fixed tcGV-expressing HEK 293T cell reacted with FlAsH. The membrane is shown in red (Lck-mScarlet-I), the nucleus in blue (DAPI), and tcGVs in green (FlAsH). All scale bars 5 μm. Three z-slices of the 3D rendering are depicted at right. The border colors of the z-slices indicate the heights of the slices within the original 3D image: z = 0.4 μm (orange, bottom), z = 2.2 μm (red, middle), z = 5.5 μm (purple, top) above the base of the cell. Corresponding notches in the 3D rendering mark the approximate heights of the z-slices. (b) – (d) Box-and-whiskers plots of the distributions of GV cluster volumes (b), distances to the nucleus (c), and distances to the membrane (d). N = 6 cells and 122 GV clusters analyzed.

After rendering the cells in 3D, we found that GVs form distinct clusters within the cell that vary considerably in size, ranging from the size of single GVs (around 0.003 μm^3^) to 20 μm^3^, with an average of 1.4 μm^3^ and a standard deviation of 2.9 μm^3^, and together occupy between 0.21% and 1.1% of the total cell volume. We then computed the distances between the GV clusters to the nuclear and plasma membranes and found that virtually all GV clusters were not in direct contact with the nucleus or the plasma membrane and remain localized to the cytosol, with the average GV cluster’s center being 2.9 ± 2.2 μm away from the nucleus and 2.6 ± 0.92 μm away from the plasma membrane (Figure 4b-d).

## CONCLUSION

In summary, our results show that intracellularly expressed AnaGVs within HEK 293T cells can be fluorescently labeled and imaged with confocal microscopy for the first time. The C-terminus of GvpA proved to be quite tolerant of mutations to cysteine, allowing for the substitution of the TC tag into GvpA without major disruption of GV expression. And, while HEK 293T cells could not synthesize GVs made entirely of tcGvpA, we found that delivering a mixture of wtGvpA and tcGvpA led to the expression of FlAsH-labelable tcGVs. We demonstrated the utility of FlAsH labeling of *in situ*-expressed GVs by studying their intracellular distribution with higher spatial precision than ever before, and found that they generally localize to the cytoplasm. We anticipate that our approach will become a tool that not only furthers the development of GV-based technologies, but also one that can be applied to the study of other genetically encoded polymeric proteinaceous structures.

## EXPERIMENTAL PROCEDURES

### Expression and screening of Serratia GV variants in E. coli on solid media

For the bacterial screens of the C-terminus of GvpA1, all mutants were cloned from the arabinose-inducible bARG_Ser_ plasmid^6^ (https://www.addgene.org/192473/) using Gibson assembly with enzyme mix (New England Biolabs, Ipswich, MA). A bacterial expression plasmid encoding GFP under the same promoter and backbone was used as a control in the same manner as the fluorescent protein controls described in Hurt et al^6^. The mutant plasmids were transformed via electroporation into Stable competent *E. coli* (New England Biolabs). Transformed *E. coli* were then patch plated onto solid inducer-free LB media containing 1.5% (w/v) agar, 1% (w/v) glucose, and 25 μg/mL chloramphenicol. Bacterial patches were made by resuspending a colony of uninduced transformed *E. coli* in 100 μL phosphate-buffered saline (PBS), then depositing 1 μL of that suspension onto both an uninduced control plate and an induced LB media plate containing 1.5% (w/v) agar, 1% (w/v) L-arabinose, 0.1% (w/v) glucose, and 25 μg/mL chloramphenicol using low-retention pipette tips. The bacterial patches were grown at 37°C for 2 days. GV expression was quantified with a ChemiDoc gel imager (Bio-Rad, Hercules, CA) by measuring the opacity of the patches. Images were processed using ImageJ (NIH, Bethesda, MD). For each GvpA1 variant (and the GFP control), four separate transformed colonies were used to make patches in case of high patch-to-patch variability. Each of these four biological replicates was patch-plated four individual times: twice onto separate induced plates, and twice more onto separate uninduced plates.

### Expression of GVs in HEK 293T cells

HEK 293T cells (American Type Culture Collection [ATCC], CLR-2316) were cultured in a humidified incubator in 0.5 mL DMEM (Corning, 50-0030PC) with 10% FBS (Takara Bio, 631368) and 1x penicillin-streptomycin in 24-well glass-bottomed No. 0 plates (Mattek, Ashland, MA, P24G-0-10-F). The plates were pre-treated with 200 μL 50 μg/mL fibronectin (Sigma-Aldrich, St. Louis, MO) in PBS at 37°C overnight before the cells were added. When the cells reached around 40% confluency, they were transfected by mixing roughly 600 ng of plasmid mixture per well with 1.6 μL Transporter 5 Transfection Reagent (Polysciences, Warrington, PA) in 60 μL 150 mM NaCl, letting the DNA complexes form for 20 minutes, then gently pipetting the solution onto the cells. Cells were transfected with a modified mARG^6^ plasmid cocktail in which each GV gene was on its own plasmid driven by a constitutive cytomegalovirus (CMV) promoter. The plasmid cocktail contained a 4:1 ratio of wtGvpA to tcGvpA, no GvpC, a 4:1 ratio of total GvpA to every other individual GV gene, and an additional plasmid encoding for Lck-mScarlet-I under a CMV promoter. Following transfection, the growth media was exchanged daily until the cells grew fully confluent (usually on day 2 post-transfection); at this point, the cells were trypsinized and re-plated onto another fibronectin pre-treated 24-well plate at a 4x dilution of their original concentration. To achieve this, the growth media was aspirated and replaced with 50 μL pre-warmed trypsin solution (Corning) per well, then the plate was incubated at 37°C for 7 minutes. The trypsin was then quenched with 550 μL DMEM per well; the contents of each well were then pipetted up and down, and 125 μL of the new suspension was transferred into 375 μL pre-warmed DMEM in a new plate for a 4x dilution. The new plate was then grown with daily media changes until the cells were roughly 60% confluent, at which point they were ready to be reacted with FlAsH.

### FlAsH reaction of cultured HEK cells and preparation for imaging

Live cultured cells were reacted with FlAsH by first washing with Hanks’ Balanced Salt Solution (HBSS, Corning 21-023-CV), then applying 250 μL of a 3 μM working solution of FlAsH-EDT_2_ (Cayman Chemical, Ann Arbor, MI) in HBSS to each well. FlAsH-EDT_2_ aliquots were prepared at 2 mM in DMSO and frozen at -80°C until use. The cells were stained with FlAsH for 30 minutes in the dark with the plate lid closed to prevent evaporation. After 30 minutes, the FlAsH working solution was removed and the cells were washed twice with a solution of 250 μM dimercaprol (also known as British Anti-Lewisite, or BAL) to reduce non-specific FlAsH binding. Pure BAL (10 molar) was purchased from Sigma-Aldrich and diluted 400x in water to make a 25 mM stock solution. The stock solution was diluted 100x in HBSS to make the BAL working solution. After the second BAL wash, the cells were fixed with 2% paraformaldehyde (Electron Microscopy Sciences, Hatfield, PA) in PBS for 20 minutes, then stained with 1 μg/mL DAPI.

### Imaging and image processing of fixed, stained cells

Cells were imaged with a Zeiss LSM 800 confocal microscope with ZEN Blue. Images were processed with the Fiji package of ImageJ. 3D renderings and measurements were performed with Imaris 10.0.1 software (Oxford Instruments, Abingdon, England, United Kingdom). In Imaris, strongly fluorescent regions in 3D space were treated as surfaces; regions of green fluorescence corresponded to tcGVs (FlAsH), red to the membrane (mScarlet-I), and blue to the nucleus (DAPI). The software was used to compute statistics relevant to these surfaces, such as the distances between them, their total volume, etc. Distances between the GV clusters and the nucleus and membrane were calculated from the distance between the estimated geometric center of each GV cluster and the nearest point on the relevant surface (nucleus/membrane).

## ACKNOWLEDGEMENTS

Imaging was performed in the Biological Imaging Facility, with the support of the Caltech Beckman Institute and the Arnold and Mabel Beckman Foundation. The authors particularly thank Dr. Giada Spigolon of the Biological Imaging Facility for her assistance with the confocal microscopy and 3D rendering of the z-stacks. This work was supported by the Chan-Zuckerberg Initiative (2020-225370 to MGS) and the National Institutes of Health (DP1 EB033154). Related work in the Shapiro lab is supported by the David and Lucile Packard Foundation. M.G.S. is an investigator of the Howard Hughes Medical Institute.

## Notes

### Competing Interest Statement

The authors have declared no competing interest.

