## Supporting Information for "Bioorthogonal labeling enables in situ fluorescence imaging of expressed gas vesicle nanostructures"

**Contents:**
**Supplementary Figures S1-S4.**

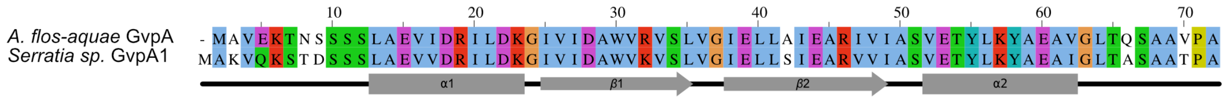


**Figure S1: Sequence alignment for *Anabaena flos-aquae* gvpA and *Serratia sp. 39006* gvpA1.** The two genes have an identity score of 80.6% (58 of 72 amino acids) and a similarity score of 91.7% (66 of 72 amino acids).

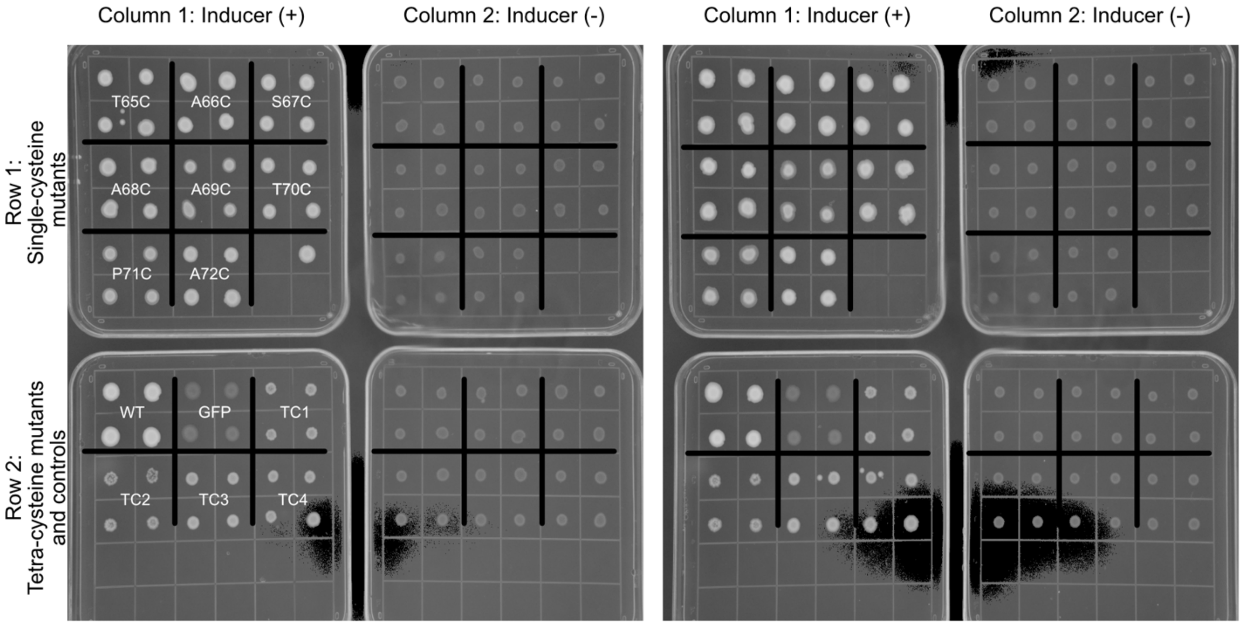


**Figure S2: Opacity screen of mutant gvpA1 plasmids in *E. coli*.** Patches of *E. coli* transformed with arabinose-inducible constructs encoding mutant gvpA1, wild type gvpA1, or GFP were grown on LB agar plates. Two images, each showing a set of four distinct plates, are shown. The patches on the four plates on the right represent duplicate technical replicates of the corresponding patches on the four plates on the left. Within each set of four plates, the two plates on the left (column 1) contain the inducer arabinose; the two plates on the right (column 2) do not. Each gvpA1 variant (and GFP) is represented by 4 patches arranged into a 2x2 grid in which each of the 4 patches originates from a different colony from the original transformation (*i.e.*, each of the 4 patches represents a distinct biological replicate). 2x2 grids of patches are labeled with their corresponding gvpA1 variant (or GFP) in the column on the left; all other columns are arranged identically.

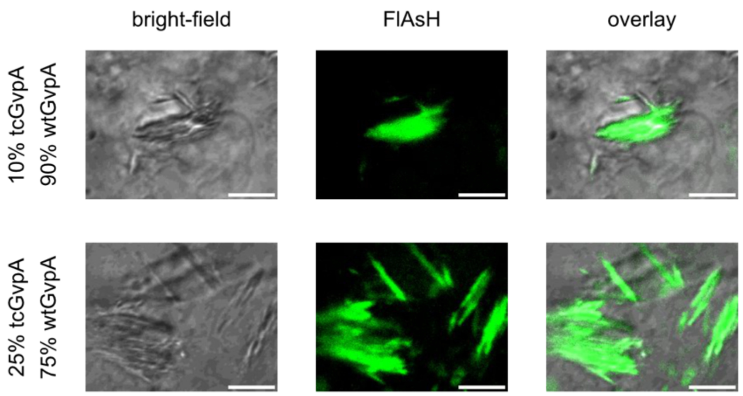


**Figure S3: Images of FlAsH-labeled GVs in cells transfected with different ratios of wtGvpA:tcGvpA.** Images of tcGV clusters in fixed HEK 293T cells transfected with 10% (top row) and 25% (bottom row) tcGvpA. GV clusters are visible under bright-field imaging (first column) and are brightly labeled with FlAsH (second column). The bright-field/FlAsH overlay (third column) demonstrates that the strongest FlAsH signal overlaps with tcGV clusters. All scale bars 5 μm.

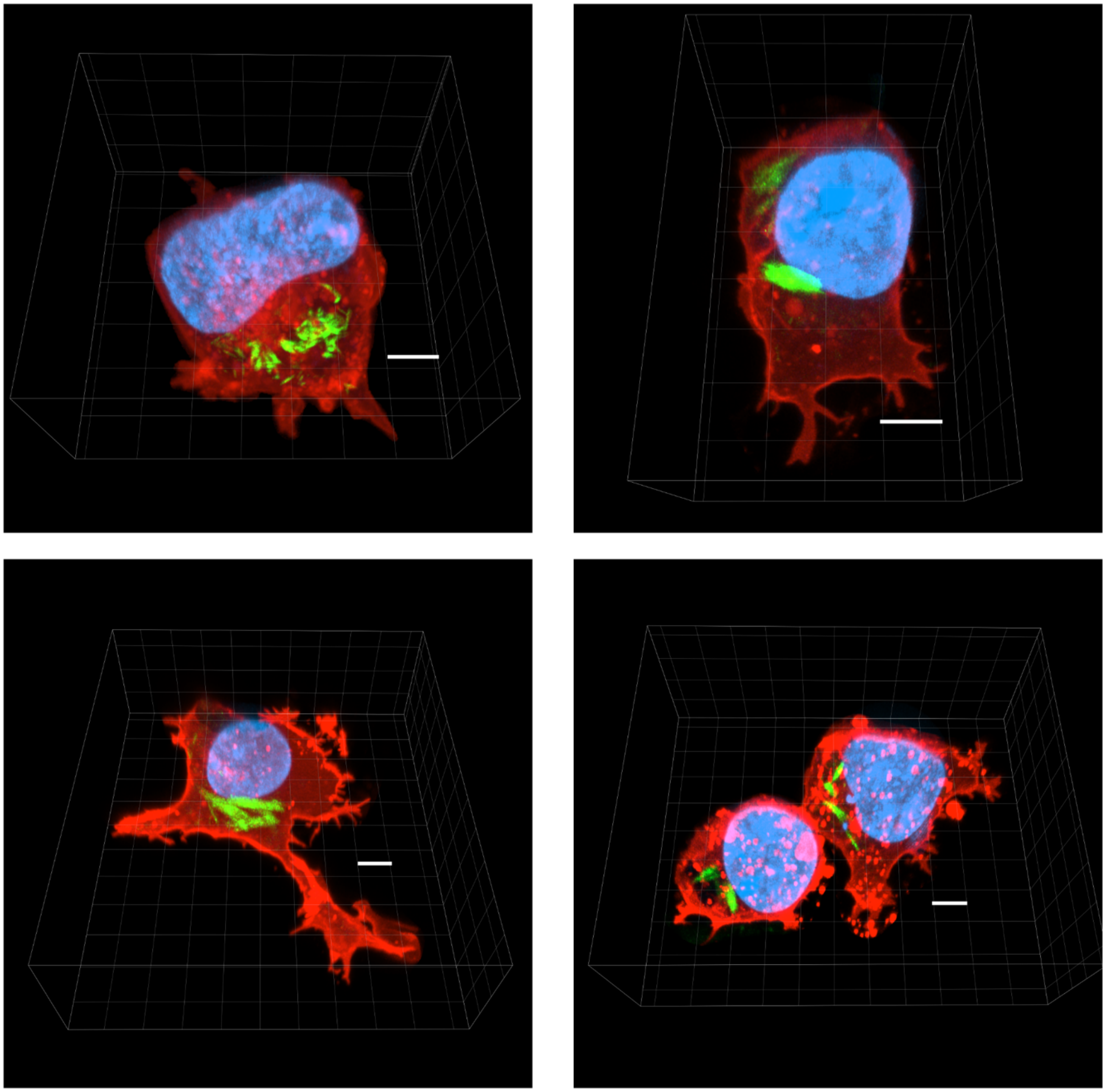

**Figure S4: 3D renderings of tcGV-expressing cells**. The cell membrane is depicted in red (Lck-mScarlet-I), tcGVs in green (FlAsH), and nucleus in blue (DAPI). All scale bars 5 μm.
